# THER: Integrative Web Tool for Tumor Hypoxia Exploration and Research

**DOI:** 10.1101/2023.11.21.568188

**Authors:** Yasi Zhang, Anqi Lin, Hong Yang, Zaoqu Liu, Quan Cheng, Jian Zhang, Peng Luo

**Author notes:** Co-corresponding Authors. Correspondence may also be addressed to: Quan Cheng, Jian Zhang, Peng Luo. Joint Authors. These authors have contributed equally to this work and share first authorship.

## Abstract

Hypoxia is an important factor in the adaptation of tumor cells to their environment, contributes to their malignant progression, and affects tumor prognosis and drug sensitivity. Although there is a wealth of transcriptomic data stored in public databases, there is a lack of web-based tools for analyzing these data to explore the link between hypoxia and the mechanisms of tumorigenesis and progression. To this end, we have developed an interactive web-based tool called THER, which is designed to help users easily identify potential targets, mechanisms of action and effective drugs for treating hypoxic tumors. THER integrates 63 transcriptomic tumor hypoxia datasets from the Gene Expression Omnibus (GEO) database, covering 3 species, 18 tumor types and 42 cell line types. This web tool provides five modules that allow users to perform differential expression analysis, expression profiling analysis, correlation analysis, enrichment analysis and drug sensitivity analysis on different datasets based on different oxygen statuses. We expect that users will be able to use the tool to identify valuable biomarkers, further reveal the molecular mechanisms of tumor hypoxia, and identify effective drugs, thus providing a scientific basis for tumor diagnosis and treatment. THER is open to all users and can be accessed without login at https://smuonco.shinyapps.io/THER/.

## Introduction

Tumor hypoxia refers to the gradual decrease in the rate of ATP production in cells or tissues when the partial pressure of oxygen drops below a critical level, and hypoxia occurs within the tumor, thus contributing to the malignant development of tumor cells to adapt to their environment(1). Several studies have shown that a series of changes induced by hypoxia play an indispensable role in tumor development; for example, hypoxia-induced acetylation of PAK1 and phosphorylation of ATG5 promote brain tumor formation(2). Under the selective pressure of hypoxia, tumors can make themselves invasive or metastatic through various mechanisms. In addition, hypoxia induces TP53 point mutations and copy number variations in many types of tumors, making them more aggressive(3). Hypoxia activates ATAD2 expression through hypoxia-inducible factor (HIF-1α), leading to increased mitochondrial reactive oxygen species (mtROS) levels, which increases the aggressiveness of lung cancer(4).

Based on the importance of hypoxia in tumorigenesis and development, hypoxia phenotypes have been used in several studies to explore their importance in tumor prognosis. For example, in cervical and head and neck cancers, patients with hypoxic tumors showed poor prognosis(5-8). Moreover, hypoxic tumors show resistance to radiotherapy and immunotherapy(8-10). However, the effect of hypoxia on the development, progression, and sensitivity to drugs may be quite different in different tumors. For example, ACHN, A2780, NT2, NCCIT, and 2102 EP cells under hypoxic conditions have decreased sensitivity to most drugs, whereas H69 and MCF7 cells have increased sensitivity to most drugs(11,12) but the mechanistic differences have not yet been clarified. Therefore, we hope to explore the mechanisms of hypoxia-induced tumorigenesis and development, identify new markers for assessing hypoxia-induced tumorigenesis and development, and identify effective drugs for treating hypoxic tumors, thus providing important guidance for the development of tumor diagnosis and treatment strategies.

Hypoxia is an important topic of interest in the field of oncology; consequently, there is an extremely rich amount of transcriptomic data stored in public databases, providing an unprecedented possibility to study the effect of hypoxia on tumorigenesis, progression and drug resistance. However, analysis of transcriptomic data to explore the link between hypoxia and the mechanisms of tumorigenesis and progression still requires a certain programming foundation, which poses a great challenge for most clinical researchers who do not have basic programming knowledge. Therefore, we developed an online web analysis tool called THER. On this platform, users can analyze the results of their research based on tumor hypoxia-associated transcriptomic datasets from the GEO database, and based on different oxygen states (hypoxia/normoxia), differential expression, expression profiling, correlation, enrichment, and drug sensitivity analyses can be easily and quickly performed. This tool can enable users to identify potential diagnostic, prognostic, predictive, and pharmacodynamic biomarkers of tumor hypoxia to further explore the molecular mechanisms involved in hypoxia-induced tumor development to provide an important scientific basis for tumor diagnosis and treatment.

## Material and Methods

The THER website (https://smuonco.shinyapps.io/THER/) allows users to individually perform differential gene expression, expression profiling, correlation, enrichment and drug sensitivity analyses. The web tool is built on the shiny package(13) for the R script (version 4.1.1, https://www.r-project.org/) and deployed based on shinyapps.io.

### Data Collection

We searched the GEO database with the keyword “hypoxia” and filtered the results. We only included (i) tumor datasets, (ii) samples treated with hypoxia or normoxia only, (iii) datasets with more than 3 samples in both the hypoxia and normoxia groups, (iv) data for *Homo sapiens*, *Mus musculus* or *Rattus norvegicus*, and (v) transcriptomic datasets. We finally collected 63 datasets as built-in datasets, including 18 tumor types and 42 cell line types.

### Data Preprocessing

#### High-Throughput Sequencing

We preprocessed the raw count data using the DESeq package(14), constructed the DESeqDataSet object by the DESeqDataSetFromMatrix function, filtered the low abundance data, performed the differential expression analysis by the DESeq function and normalized the matrix of expression values by the rlog function.

#### Affymetrix Arrays

We used the affy package(15) to preprocess the raw data of the HGU95, HGU133 and MGU74 series, read the raw data with the ReadAffy function, and perform background correction, normalization and expression calculation with the rma function on the raw data. We used the oligo package(16) to preprocess the raw data of other series, read the raw data with the read. celfiles function, and perform background correction, normalization and expression value calculation with the rma function on the raw data.

#### Illumina BeadArray

We used the lumiR.batch function and the lumiExpresso function in the lumi package(17) to read, background correct and quantile standardize the raw data.

#### Agilent arrays

We used the backgroundCorrect function and normalizeBetweenArrays function in the limma package(18) to background correct and quantile standardize the raw data, respectively.

### Analysis Modules

#### Data

We used the DT package (https://rstudio.github.io/DT/shiny.html) to present specific information about the 63 built-in datasets and set the table styles with css. Using the shinydashboard package, the number of datasets, hypoxia sample sizes, and normoxia sample sizes were presented in the table.

#### Differential Expression Analysis

We used the limma package for gene expression differential analysis, and the p value obtained from the t test was linked to the mean expression of the hypoxia group minus the mean expression of the normoxia group (i.e., log2FC) to determine whether the genes were significantly different genes. The p value cutoff was set to 0.05 by default, and the |log2FC| value cutoff was set to 1 by default. We then displayed the results of the differential expression analyses in visual charts using the ggplot2 (version 3.3.5, https://cran.r-project.org/web/packages/ggplot2/index.html) and ggrepel packages for volcano plots, the pheatmap package(19) for heatmaps, and the DT package for tables.

#### Expression Analysis

The p value indicating the significance of differentially expressed genes was calculated by the Mann□Whitney U test. We presented the results of the expression profiles in visual charts, using the ggplot2 package to draw boxplots and barplots, using the ggpubr package to display the p values in graphs, and using the DT package to generate tables.

#### Correlation Analysis

We downloaded GO terms, KEGG pathways, Reatome pathways, and Wikipathways from the Molecular Signatures Database (MsigDB)(20). The relative expression activity of the pathways was calculated by including all genes in the selected dataset using the ssgsea method with the gsva function in the GSVA package(21). Our correlation analyses use the Spearman method by default and allow the user to select the Pearson method. We then presented the results of the correlation analysis in visual charts, using the ggplot2 package to plot scatter plots and the ggpubr package(22) to display the correlation coefficients and p values in the plots, using the corrplot package(23) to generate correlation heatmaps, and using the circlize package(24) to establish chord diagrams.

#### Enrichment Analysis

We used the Over Representative Analysis (ORA) and Gene Set Enrichment Analysis (GSEA) algorithms in the clusterProfilter(25) and ReactomePA(26) packages as well as the Gene Set Variation Analysis (GSVA) algorithm in the GSVA package to perform enrichment analysis based on four annotation databases: GO terms, KEGG pathways, Wikipathways, and Reatome pathways. In the GSEA and GSVA algorithms, we included all genes in the selected datasets, while in the ORA algorithm, we only included significantly differentially expressed genes in the selected datasets. The p value and q-value intercept values were set to 0.05 in both the GSEA and ORA algorithms, while the GSVA algorithm set |log2FC| to 1 and the p value to 0.05 in both algorithms. Subsequently, we presented the results of the enrichment analyses in visual charts, using the enrichplot package(27) to plot dotplots, barplots and GSEA plots, the ggplot2 package to plot ridgeplots, the pheatmap package to plot heatmaps, and the DT package to produce tables.

#### Drug Sensitivity Analysis

We used the pRRophetic package(28) to calculate the concentration of drug required to inhibit half of the tumor cells (IC50). The drug parameter of the pRRopheticPredict function has 138 drug names, and calculating these 138 drugs gives us. The result is the IC50 after processing with log. Subsequently, we presented the results of the drug sensitivity analysis in visual charts, using the ggplot2 package for boxplots, the pheatmap package for heatmaps, and the DT package for tables.

### Example

We checked the number of samples of cell line types included in this web tool at the bottom of the Home page and selected the cell line with the largest number of samples to be studied. Using the "DEG" module, one of the datasets from the cell line under investigation was selected for differential analysis, and the top five significantly differentially expressed genes in descending order of |log2FC| values were screened. (|log2FC| > 1; p value ≤ 0.5) as candidate marker genes. Subsequently, we searched PubMed for these candidate marker genes and found that four of them had been reported as differential hypoxia genes in the cell lines under investigation, and the unreported genes were used as differential hypoxia genes in the cell lines under investigation. To further verify the reliability of the target genes as candidate marker genes, the "DEG" module was used again, and it was found that the candidate marker genes were significantly upregulated in six of the seven MCF7 cell line datasets. At the same time, to further investigate the role of this gene in the adaptation of tumors to hypoxic environments, we used the "Enrichment" module and performed enrichment analysis based on the ORA method. In addition, we used the "Correlation" module to perform correlation analysis based on the Spearman method to explore the correlation between the expression of the candidate marker genes and the expression of the pathway-related genes to verify the mechanism of the candidate marker genes.

## Results

THER preprocesses 63 hypoxia-associated transcriptome datasets in advance so that users can perform analysis and visualization easily and quickly. THER contains five analysis modules, namely, the differential expression analysis module, expression profiling module, correlation analysis module, enrichment analysis module, and drug sensitivity analysis module (Figure 1). Under the five analysis modules, there are 17 subanalysis modules, and a description button is designed in the upper left corner of the output page of each subanalysis module, which is designed to help the user quickly understand the operation method and meaning of each subanalysis module. Users can use these analysis modules to autonomously explore various features in tumor samples under hypoxia/normoxia conditions to investigate the changes that occur in tumors in hypoxic environments. The analyses are available for download, allowing users to export spreadsheets in csv format and high-resolution images in both png and pdf formats and to customize the length and width of the images.

**Figure 1:**
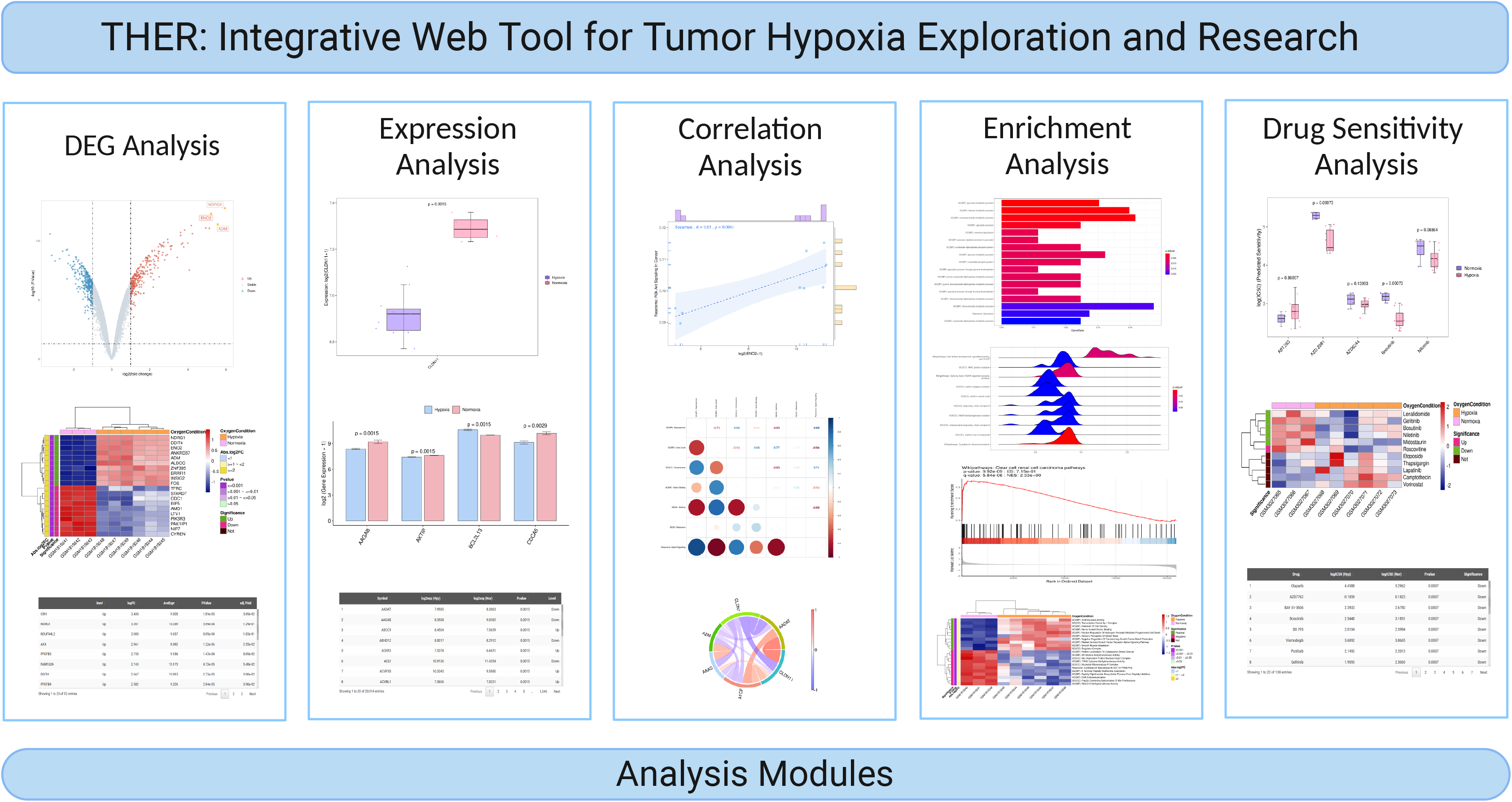
Analysis modules of THER. THER provides users with five analysis modules, including a differential gene expression analysis module, an expression profiling module, a correlation analysis module, an enrichment analysis module, and a drug sensitivity analysis module.

### Data

Below the Data module, general information about the 63 datasets included in THER is shown,and more detailed information about the datasets is available to the user by clicking on the plus button on the left side of each column in the table; thus, the user can quickly locate the dataset of interest and jump directly to the analysis module of the corresponding dataset by clicking on the black jump button in the first column of the table. Additionally, this module provides users with a search bar for filtering datasets of interest based on cancer types and cell line types and displays the number of eligible datasets, the number of samples with hypoxia, and the number of samples with normoxia at the top of the page.

### Differential Expression Analysis

The DEG analysis module consists of three submodules, volcano, heatmap and table (limma), which allow the user to compare the differences in gene expression between the normoxia and hypoxia groups and to present the results of the limma differential expression analysis according to oxygen status in different formats (Figure 2).

**Figure 2:**
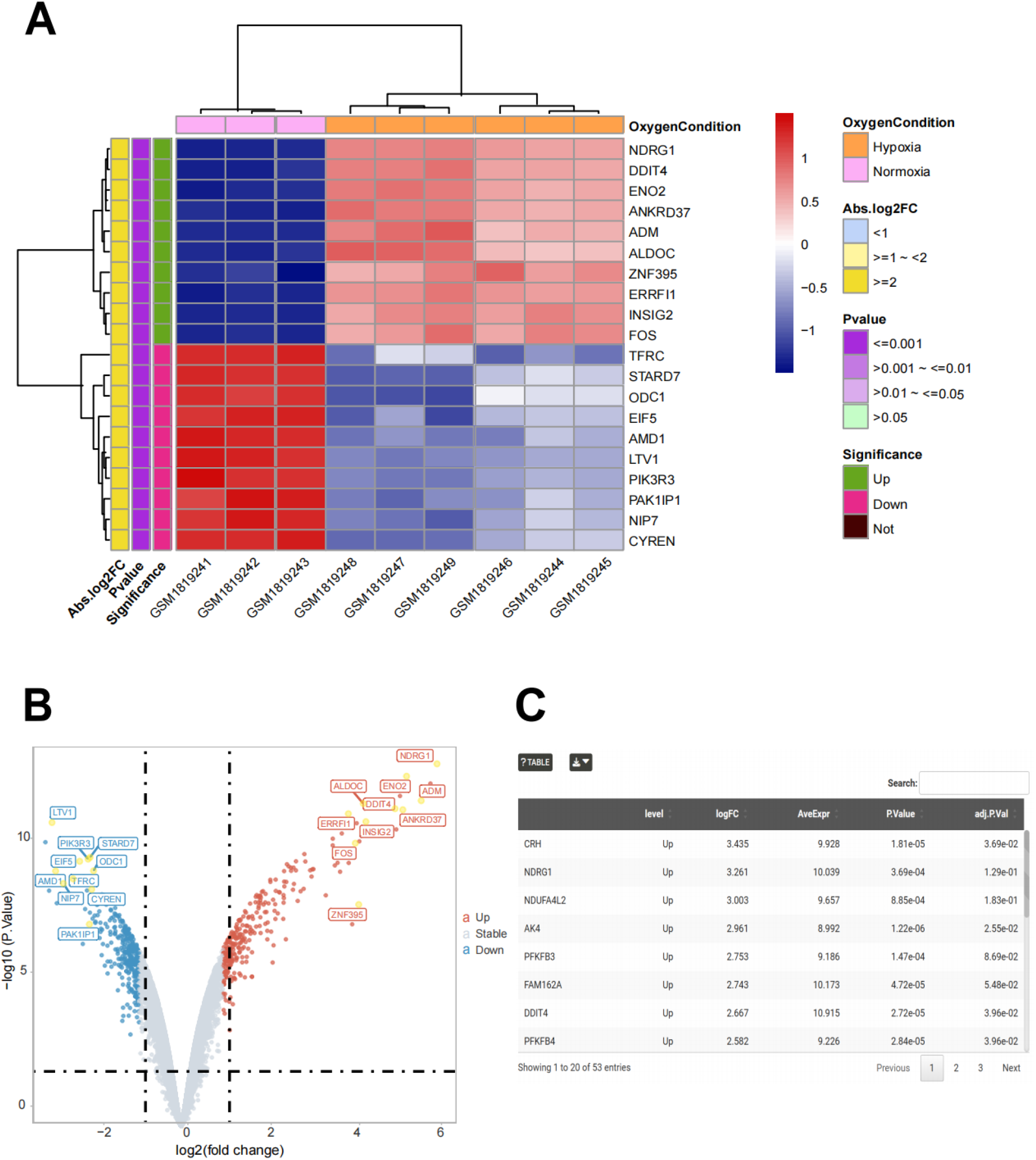
Visualization of the results of differential expression analysis obtained from limma analysis, showing whether specific genes were significantly upregulated or downregulated between the hypoxia and normoxia groups in different visualizations. (A) In the heatmap, the z-score-transformed gene expression values for each sample are shown in the cells, and the significance and trend of significance of the genes are shown in the right-hand labels. (B) In the volcano plot, red dots represent significantly upregulated genes, blue dots represent significantly downregulated genes, and gray dots represent nonsignificant genes. (C) Summary table of the results of differential expression analysis.

#### Volcano

The submodule volcano allows the user to visualize volcano plots indicating whether the expression of genes are significantly upregulated, significantly downregulated or not significantly regulated in the hypoxia group relative to the normoxia group (Figure 2B). Users are free to select the dataset of interest and can choose to display custom genes or the top genes in descending order by |log2FC| value, i.e., the most upregulated and downregulated genes, for visualization. The user can also adjust the plot by selecting the p-value cutoff, the |log2FC| cutoff and the color scheme of the volcano plots in the panel that expands when the “Extra Parameters” button is clicked.

#### Heatmap

The submodule heatmap allows users to visualize heatmaps illustrating the expression levels of genes in different samples and the results of significant differences in different oxygen states (Figure 2A). Users are free to select the dataset of interest and can choose to display custom genes or visualize the top genes in descending order by |log2FC| value, i.e., the most upregulated and downregulated genes. Users can also adjust the plot by choosing whether to normalize by rows or columns, whether to show values in heatmaps, whether to cluster by columns, and the color scheme of the heatmaps in the panel that expands when the “Extra Parameters” button is clicked.

#### Table (limma)

The table submodule allows users to view the results of the differential expression analysis in tabular formats (Figure 2C), including the level of significant difference, |log2FC| value, mean expression value or count mean, p value and adjusted p value. This module supports not only the user’s choice to display all genes but also the user’s choice to display only significantly differentially expressed genes; thus, the user can quickly filter the results.

### Expression Analysis

The expression module consists of three submodules, boxplot, barplot and table (wilcox-test), which allow users to compare the expression differences of single or multiple genes between the normoxia and hypoxia groups, to present the expression values of the genes in different oxygen states in different formats and to determine whether the genes being investigated are significantly differentially expressed genes between oxygen states by using a Mann□Whitney U test (Figure 3). The Mann□Whitney U test was used to determine whether the genes explored were significantly differentially expressed genes between oxygen states (Figure 3).

**Figure 3:**
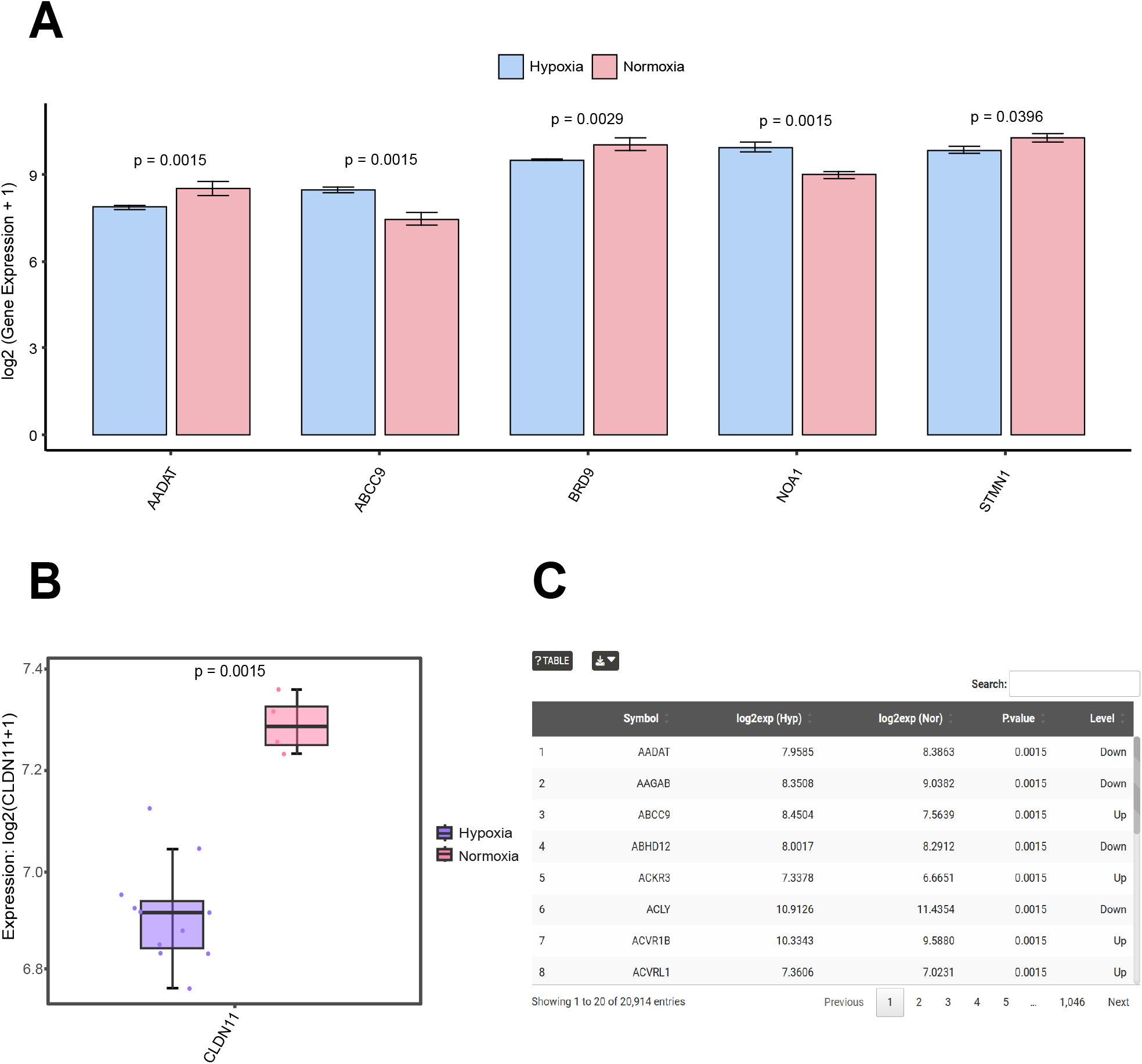
Graphical representation of the visualization results of expression profiling, showing the expression values of specific genes in the hypoxia and normoxia groups in different visualization forms, and p values for the differences in expression of genes between the groups of different oxygen states were obtained by the Mann ‒ Whitney U test. (A) Visualization of the distribution of expression values of multiple genes among different oxygen status groups in the form of a barplot. (B) Visualization of the distribution of expression values of individual genes among different oxygen status groups in the form of a boxplot. (C) Summary table of expression profiling results.

#### Boxplot

In the boxplot submodule, users can select a single gene of interest and show its differential expression in the hypoxia and normoxia groups as boxplots (Figure 3B). Users can also adjust the plot by selecting the p value presentation format and the color scheme of the boxplots in the panel that expands when the “Extra Parameters” button is clicked.

#### Barplot

The Barplot submodule allows the user to select two to nine genes of interest and present the differential expression of the selected genes between groups in the form of barplots (Figure 3A). Users can also adjust the image by selecting the p value presentation and the color scheme of the barplots in the panel that expands when the “Extra Parameters” button is clicked.

#### Table (Wilcox Test)

In this submodule, users can view the expression values, p values, whether the genes are significantly differentially expressed, and the trend of significantly differentially expressed genes in the hypoxia and normoxia groups after log2 conversion in tabular formats (Figure 3C).

### Correlation Analysis

The correlation analysis module contains three submodules, scatter, correlation heatmap and chord diagram, which allow the user to freely compare gene-to-gene, gene-to-pathway, and pathway-to-pathway correlations to explore possible connections (Figure 4).

**Figure 4:**
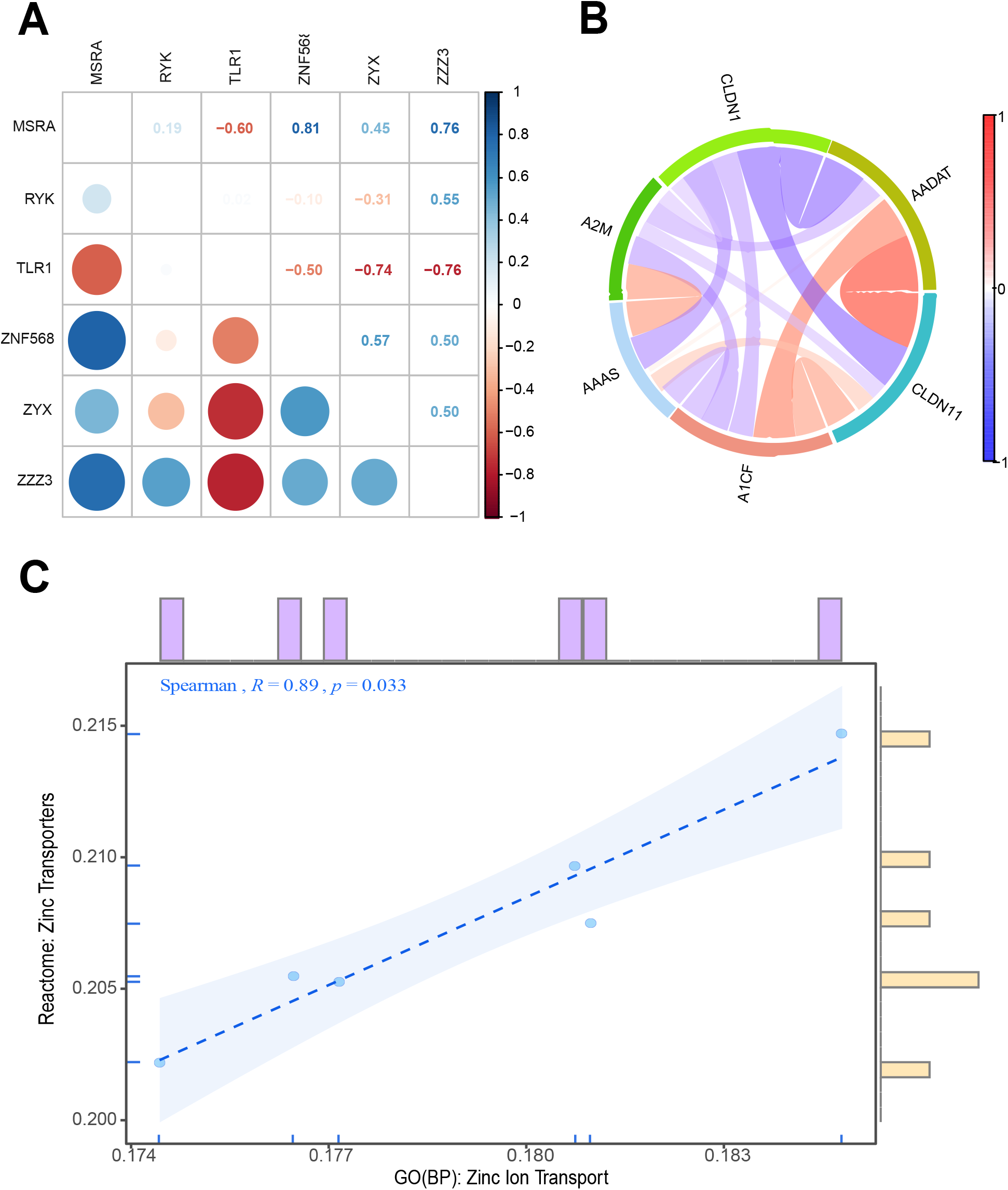
Graphical representation of the visualization results of the correlation analysis, showing the results of the correlation analysis in different visualizations. (A) In the correlation heatmap, blue indicates a positive correlation, and red indicates a negative correlation, with darker colors representing stronger correlations. (B) In the chord plot, red lines represent positive correlations, blue lines represent negative correlations, and darker colors or thicker lines represent stronger correlations. (C) In the scatter plot, the correlation coefficient R and p value are shown in the upper left corner.

#### Scatter

In the scatter submodule, users can freely choose to explore the correlation between the expression of two different genes, the relative expression activity of a gene and a pathway, or the relative expression activity of two different pathways and present the visualization results in the form of scatter plots (Figure 4C). Users can also adjust the plot by selecting the correlation analysis method, the type of edge plot and the color scheme of the scatter plot in the panel that expands after clicking the “Extra Parameters” button.

#### Correlation Heatmap

The correlation heatmap submodule allows users to freely choose to explore the correlation between the expression of multiple different genes, the expression of multiple genes with the relative expression activity of multiple pathways, or the relative expression activity of different pathways with a correlation heatmap. The results are presented graphically (Figure 4A). The user can also adjust the plot by selecting the correlation analysis method and the visualization shape of the correlation heatmaps in the panel that expands after clicking on the “Extra Parameters” button.

#### Chord Diagram

In the chord diagram submodule, users can freely choose to explore the correlation between the expression of multiple different genes, the expression of multiple genes and the relative expression activity of multiple pathways, or the relative expression activity of different pathways and present the visualization results in the form of chord diagrams (Figure 4B). Users can also adjust the plot by selecting the correlation analysis method and the transparency of the central lines of the chord diagrams in the panel that expands after clicking the “Extra Parameters” button.

### Enrichment Analysis

The Enrichment Analysis module consists of five submodules: dotplot/barplot, ridgeplot, GSEA plot, heatmap (GSVA), and table, which present the results of differentially enriched pathways in different formats (Figure 5). This module helps users correlate genes with functions and provides an opportunity to discover potential biological pathways related to tumor hypoxia.

**Figure 5:**
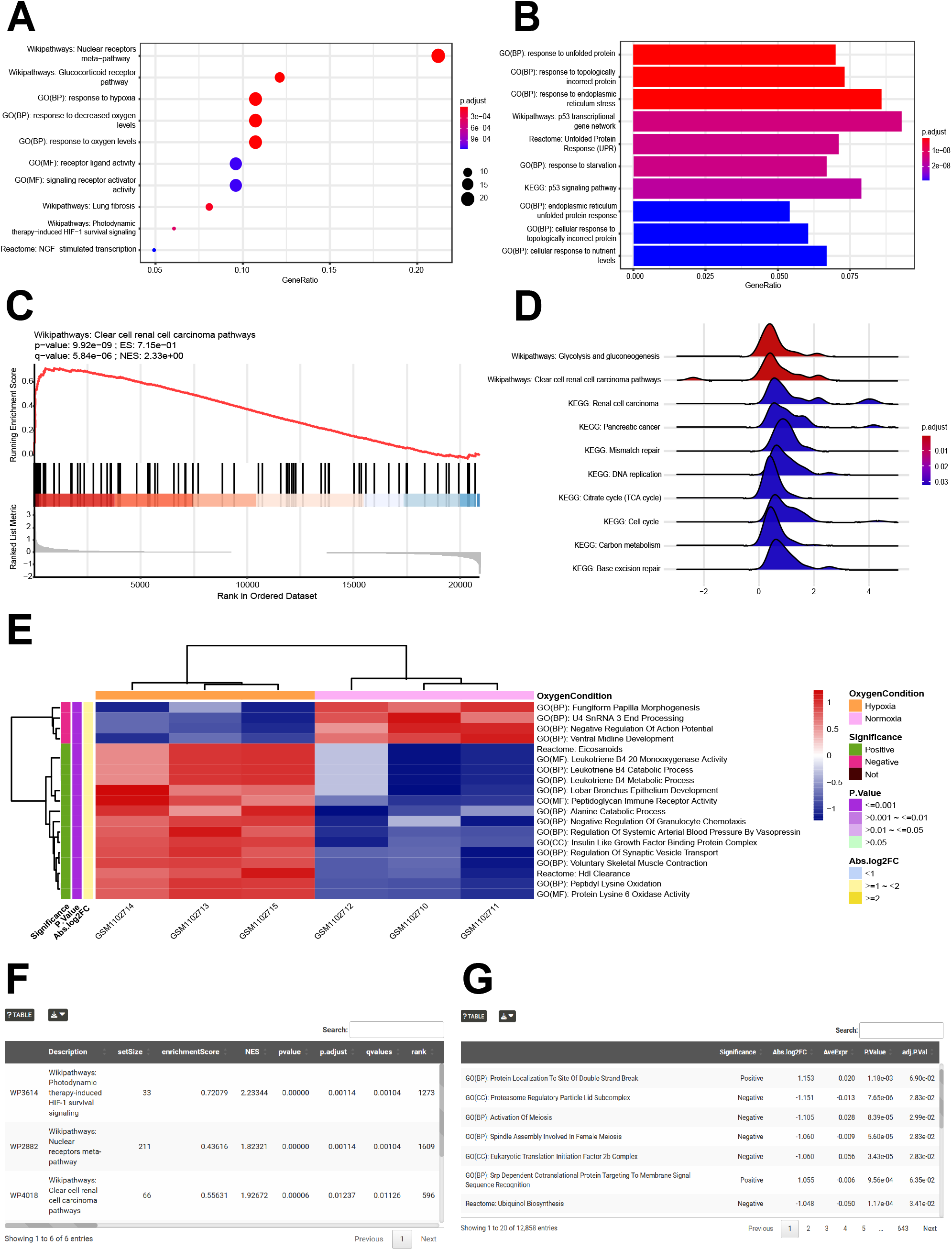
Graphical representation of the visualization results of enrichment analysis for specific pathways in different visualization forms. (A) Visualization of p.adjust values, GeneRatio and Count values for specific pathways in the form of a dotplot. (B) Visualization of the p.adjust value and GeneRatio of specific pathways in the form of a barplot. (C) In the GSEA plot, the peaks of the red curves are the enrichment scores for the set of genes, with positive values indicating top enrichment, genes to the left of the peaks are core genes, and the opposite is true for negative values. The black line indicates where the genes in the sorted table of expressed genes are located in the currently analyzed set of functionally annotated genes. The red and blue heatmaps are expression abundance rankings, where a darker red line indicates a larger logFC for the gene at that position, and a darker blue line indicates a smaller logFC. (D) Visualization of the p.adjust value and the expression distribution of core-enriched genes of specific pathways in the form of a ridgeplot. (E) In the heatmap, the GSVA scores of pathways transformed by z scores for each sample are shown in the cells, and the significance and trend of significance of the pathways are shown in the right-hand labels. (F) Summary table of enrichment analysis results by the ORA enrichment method. (G) Summary table of enrichment analysis results by the GSEA enrichment method. (H) Summary table of enrichment analysis results by the GSVA enrichment method.

#### Dotplot/Barplot

This submodule allows users to select either ORA or GSEA methods. Users are free to select the dataset of interest and can choose to visualize either the top pathways in ascending order by p value or custom pathways. The visualization of differentially enriched pathways in different oxygen states obtained by ORA or GSEA methods is shown as dotplots (Figure 5A). The visualization of the differentially enriched pathways obtained by the GSEA method is shown as barplots (Figure 5B). Users can also adjust the plot by selecting the variable represented by the X-axis and the variable represented by the color in the panel that expands after clicking the “Extra Parameters” button.

#### Ridgeplot

The ridgeplot submodule uses the GSEA enrichment analysis method to show the visualization results of differential pathways in different oxygen states in the form of ridgeplots (Figure 5D). Users are free to select the dataset of interest and can choose the top pathways in ascending order by p value or custom pathways for visualization. The user can also adjust the plot by selecting the variable represented by the color of the fill in the ridgeplot and the transparency of the ridgeplot in the panel that expands after clicking the “Extra Parameters” button.

#### GSEA Plot

The GSEA plot submodule uses the GSEA method to show the visualization results of differential pathways in different oxygen states in the form of GSEA plots (Figure 5C). Users are free to select the dataset of interest and the pathway of interest for visualization. Users can also adjust the plot by selecting the form and color of the enrichment curves in the GSEA plot in the panel that expands after clicking the “Extra Parameters” button.

#### Heatmap (GSVA)

This submodule uses the GSVA enrichment analysis method to present the results in the form of heatmaps (Figure 5E), visualizing the GSVA score levels of the pathways in different samples as well as the results of significant differences in different oxygen states. Users are free to select the dataset of interest and can choose the top pathways in descending order of |log2FC| values or customized pathways for visualization. Users can also adjust the plot by selecting whether to normalize by row or column, display values in heatmaps, cluster by column and change the color scheme of the heatmaps in the panel that expands when the “Extra Parameters” button is clicked.

#### Table

The table submodule allows users to view the enrichment analysis results of the dataset of interest performed by the ORA, GSEA and GSVA enrichment methods in tabular form (Figure 5F-G).

### Drug Sensitivity Analysis

The drug sensitivity analysis module contains three submodules, heatmap, boxplot and table, which allow users to compare the differences in drug sensitivity between samples in different oxygen states. The three submodules present the results of the drug sensitivity analyses in different formats and use the Mann□Whitney U test to determine whether or not the differences in the logIC50 values for the drugs presented are significant between samples in the hypoxia and normoxia groups (Figure 6).

**Figure 6:**
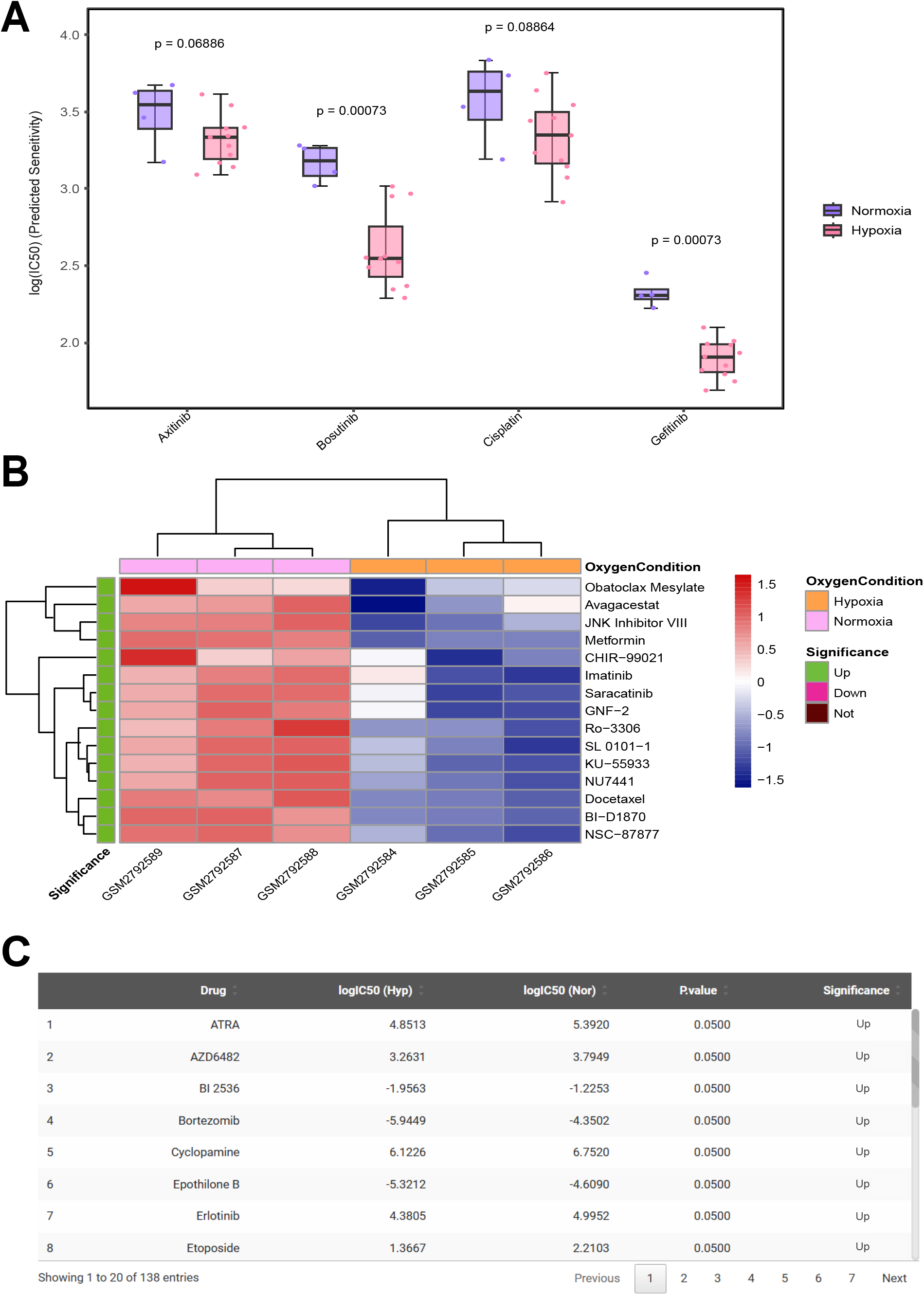
Graphical representation of the visualization results of the drug sensitivity analysis. The logIC50 values of specific drugs in the hypoxia and normoxia groups are shown in different visualizations, and the p values of the differences in drug expression between groups of different oxygen statuses were obtained by the Mann□Whitney U test. (A) Visualization of the distribution of logIC50 values for multiple drugs among different oxygen status groups in the form of a boxplot. (B) In the heatmap, the logIC50 values of the drugs transformed by z scores for each sample are shown in the cells, and the significance of the genes and the trend of significance are shown in the right-hand labels. (C) Summary table of the results of drug sensitivity analyses.

#### Heatmap

This submodule allows users to visualize the level of semi-inhibitory concentration of the drugs in different samples in the form of heatmaps (Figure 6B) and significantly different results in different oxygen states. Users are free to select the dataset of interest and can choose to visualize the customized drugs or the top drugs in descending order of |log2FC| values, i.e., the drugs with the most upward and downward adjustments. Users can also adjust the plot by selecting whether to normalize by row or column, show values in the heatmap, cluster by column and change the color scheme of the heatmap in the panel that expands after clicking on the “Extra Parameters” button.

#### Boxplot

In the boxplot submodule, users can select two to nine drugs of interest and present the difference in logIC50 values between the hypoxic and normoxic groups in the form of a box plot (Figure 6A). Users can also adjust the plot by selecting the p value presentation format and the color scheme of the boxplots in the panel that expands when the “Extra Parameters” button is clicked.

#### Table

Users can view the logIC50 values, p values, the level of significant differences, and the trend of significant differences for drugs in the hypoxia and normoxia groups in a tabular format in this submodule (Figure 6C).

### Example

We chose the cell line MCF7 (Figure 7A), which contains the largest number of samples in this web tool, for multimodule exploration. One of the datasets of the MCF7 cell line, GSE70805, was selected for differential expression analysis to screen the top five most significantly differentially expressed genes (Figure 7B). All except ENO2 have been reported as differential hypoxia genes in MCF7 cells(29-32) (Figure 7C). Moreover, ENO2 has been reported to be significantly upregulated in hypoxia-associated solid tumors, including liver cancer(33), laryngeal cancer(34), gastric cancer(35), glioblastoma(36) and melanoma(37), and could be a candidate target. However, there is no report on the changes in and role of ENO2 in hypoxia-treated MCF7 breast cancer cells. Based on the above findings, we suggest that ENO2 could be a candidate marker gene for hypoxia-treated MCF7 cells. Moreover, ENO2 was found to be significantly upregulated in six of the seven MCF7 cell line datasets (Figure 7C, Figure S1), which further validated the reliability of ENO2 as a candidate marker gene. ENO2 has been reported to function in hypoxia-associated tumors through the glycolytic pathway(33,38). Similarly, enrichment analysis showed that multiple glycolytic pathways were differential pathways in MCF7 cells (Figure 7E), and all of these pathways were involved in ENO2 (Figure 7F). To further validate the pathway of ENO2 in tumor adaptation to a hypoxic environment, we performed Spearman correlation analysis and found a significant positive correlation between ENO2 and glycolysis-related genes (Figure 7D, Figure S2). In addition, pathway enrichment analysis revealed that ENO2 plays a role in the dysregulation of nucleotide metabolism (Figure S3). This may reveal potential pathways for ENO2 to promote the rapid adaptation of tumor cells to hypoxic environments through its involvement in glycolytic reprogramming and the dysregulation of nucleotide metabolism.

**Figure 7:**
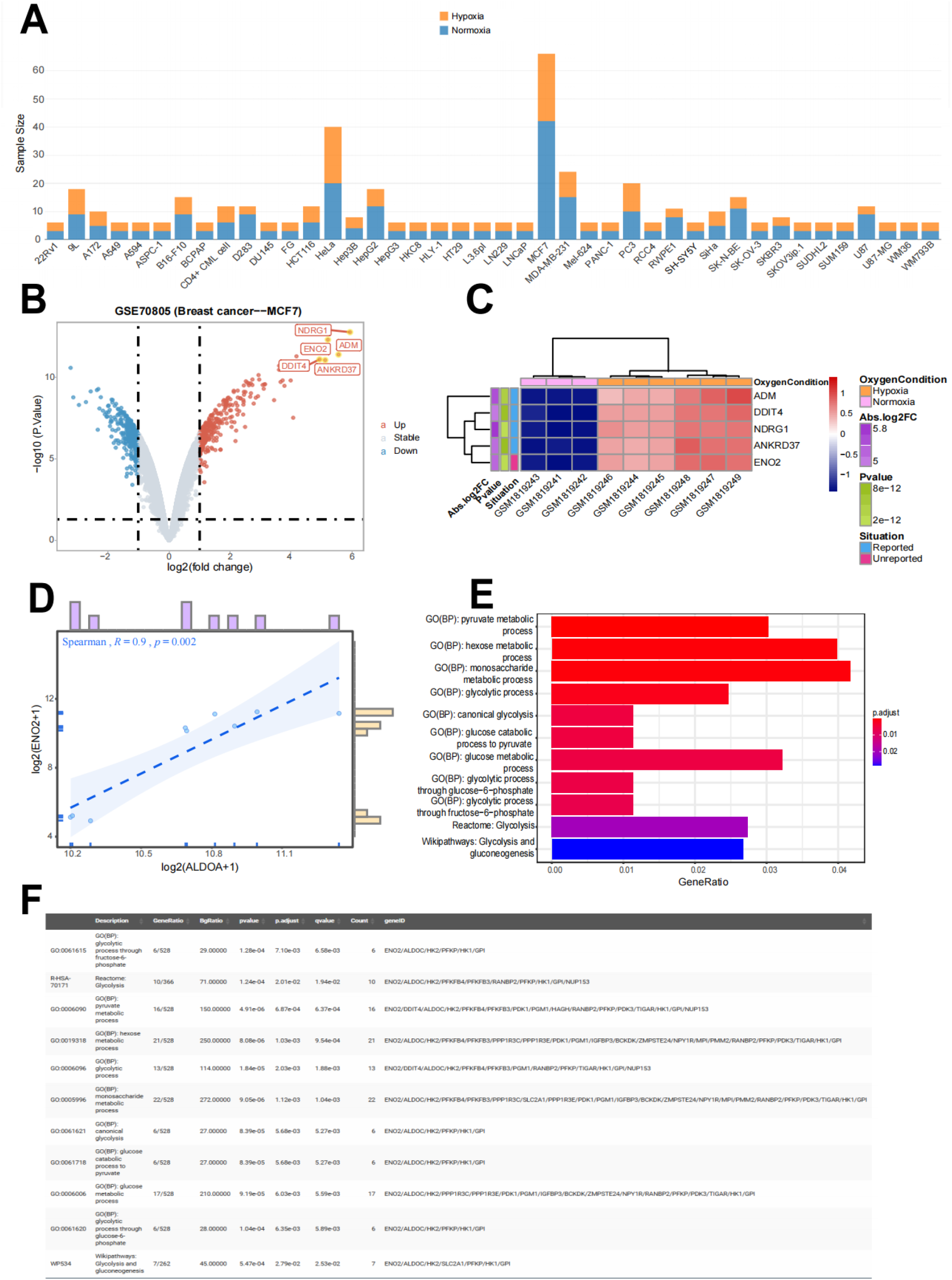
Reliability of ENO2 as a new marker for hypoxic MCF7 cells and its mechanism of action. (A) Stacked histograms showing the total sample sizes, hypoxia sample sizes and normoxia sample sizes of the cell lines included in THER’s built-in datasets. (B) Significantly differentially expressed genes of GSE70805 (breast cancer--MCF7) obtained using limma difference analysis based on different oxygen statuses. (C) Reported status of the top five most significantly differentially expressed genes of GSE70805 (breast cancer--MCF7). (D) Scatter plot showing the correlation between ENO2 and another glycolysis-related gene. (E) Demonstration of differential glycolytic pathways in the form of a barplot. (F) Demonstration of core genes of differential glycolytic pathways in the form of a table. (H) Demonstration of differential nucleotide metabolism pathways in the form of a barplot.

## Discussion

Hypoxia drives tumorigenesis and makes tumors more likely to acquire invasiveness, and it is particularly important to clarify the molecular mechanisms of hypoxia in tumorigenesis and development. In addition, identifying effective drugs during tumor hypoxia and screening for potential diagnostic, prognostic, predictive, and pharmacodynamic markers has important research and clinical implications. To address this need, we designed and successfully developed an online web tool called THER for comprehensive analysis and visualization of hypoxia-related data. The built-in datasets of the platform are 63 tumor hypoxia-related transcriptome datasets derived from the GEO database, covering 3 species (*Homo sapiens*, *Mus musculus* and *Rattus norvegicus*), 18 tumor types, and 42 cell line types.

THER is a feature-rich web tool for tumor hypoxia that provides users with five analysis modules. First, in the DEG module and expression module, users can screen for potential biomarkers. Next, in the correlation module and enrichment module, users can explore in depth the molecular mechanisms of hypoxia-induced tumor development. Subsequently, users can identify effective drugs against hypoxic tumors in the drug sensitivity module. In conclusion, this web-based tool explicitly assists clinical doctors and researchers who may not be adept in programming to easily identify potential targets, function mechanisms and potentially effective drugs against hypoxic tumors that may lead to discovering new targets and therapeutic approaches against these tumors.

THER serves as an online web tool that can provide users with an opportunity to explore potential tumor hypoxia-associated biomarkers and investigate their underlying mechanisms. For example, ENO2 may promote tumor progression toward malignancy through the glycolytic pathway and may act synergistically with other glycolysis-related genes, which is consistent with existing reports(11,33). ENO2 encodes a glycolytic enzyme involved in the energy release phase of glycolysis(39,40). Under hypoxic conditions, breast cancer cells exhibit an enhanced glycolytic phenotype, resulting in upregulation of the expression of genes encoding glycolytic enzymes, including ENO2, and increased glycolytic energy production(38). Moreover, silencing ENO2 shows synergistic effects with anti-glycolytic drugs(41). Furthermore, in breast cancer cells, simultaneous inhibition of ENO2 and the activity of another glycolytic gene, ALDOC, significantly suppresses lactate production and reduces the adaptability of tumor cells to the hostile environment(11). This example demonstrates the significant value of the THER web tool in studying tumor hypoxia-related biomarkers and mechanisms to discover potential targets for tumor therapy.

Notably, THER only explores the effect of direct oxygen status on tumor cells and does not yet address factors that indirectly affect oxygen status (e.g., different altitude levels and HIF-1α expression levels). In addition, THER has some shortcomings: 1) The sample size of the dataset is limited, so there is a risk that the error distribution may deviate from the normal distribution(42). 2) The number of datasets included for some cancer types or cell lines is small. Nevertheless, the datasets will be continuously updated in THER. 3) Many studies have already demonstrated that the effect of hypoxia on the tumor microenvironment and the interaction between hypoxia and immune cell infiltration is very complex(43-45). Therefore, the THER immune infiltration analysis module will be updated.

## Conclusion

In this study, an online web tool called THER was successfully designed and developed to comprehensively analyze and visualize hypoxia-related data. The web tool has 63 built-in transcriptomic datasets for 18 cancer types derived from the GEO database and allows users to individually and systematically perform differential expression analysis, expression profiling analysis, correlation analysis, enrichment analysis, and drug sensitivity analysis. We demonstrate, through an example, the value of the THER web tool in studying tumor hypoxia-related biomarkers and mechanisms to identify potential targets for tumor therapy. This web tool explicitly helps clinicians and researchers with no programming background to easily explore the changes that occur in tumors in hypoxic environments to delve into the potential targets, mechanisms of action and potentially effective drugs for treating hypoxic tumors, thus providing important guidance for the development of tumor diagnosis and treatment strategies.

## Data Availability

The THER website is available as a web-accessible open resource at https://smuonco.shinyapps.io/THER/. This website is built on R Studio (https://www.r-project.org/) and the shiny R package (https://github.com/rstudio/shiny). The raw data for THER’s built-in datasets were derived from the Gene Expression Omnibus database (https://www.ncbi.nlm.nih.gov/gds). All data analysed in this study are available upon reasonable request from the corresponding authors.

## Funding

None

## Figure Legend

Supplementary Figure 1: (A-D) Significance and trend of significance of ENO2 in the MCF7 cell line datasets (GSE29406, GSE47533, GSE111246, GSE111259) obtained using limma difference analysis.

Supplementary Figure 2: (A-D) Scatter plot showing the correlation between ENO2 and glycolysis-related genes (GPI, HK2, ALDOC, PFKFB3).

Supplementary Figure 3: (A) Demonstration of differential nucleotide metabolism pathways in the form of a barplot. (B) Demonstration of core genes of differential nucleotide metabolism pathways in the form of a table.

## Notes

### Competing Interest Statement

The authors have declared no competing interest.

