## Supplementary figures and images for "THER: Integrative Web Tool for Tumor Hypoxia Exploration and Research"

### Figure S1

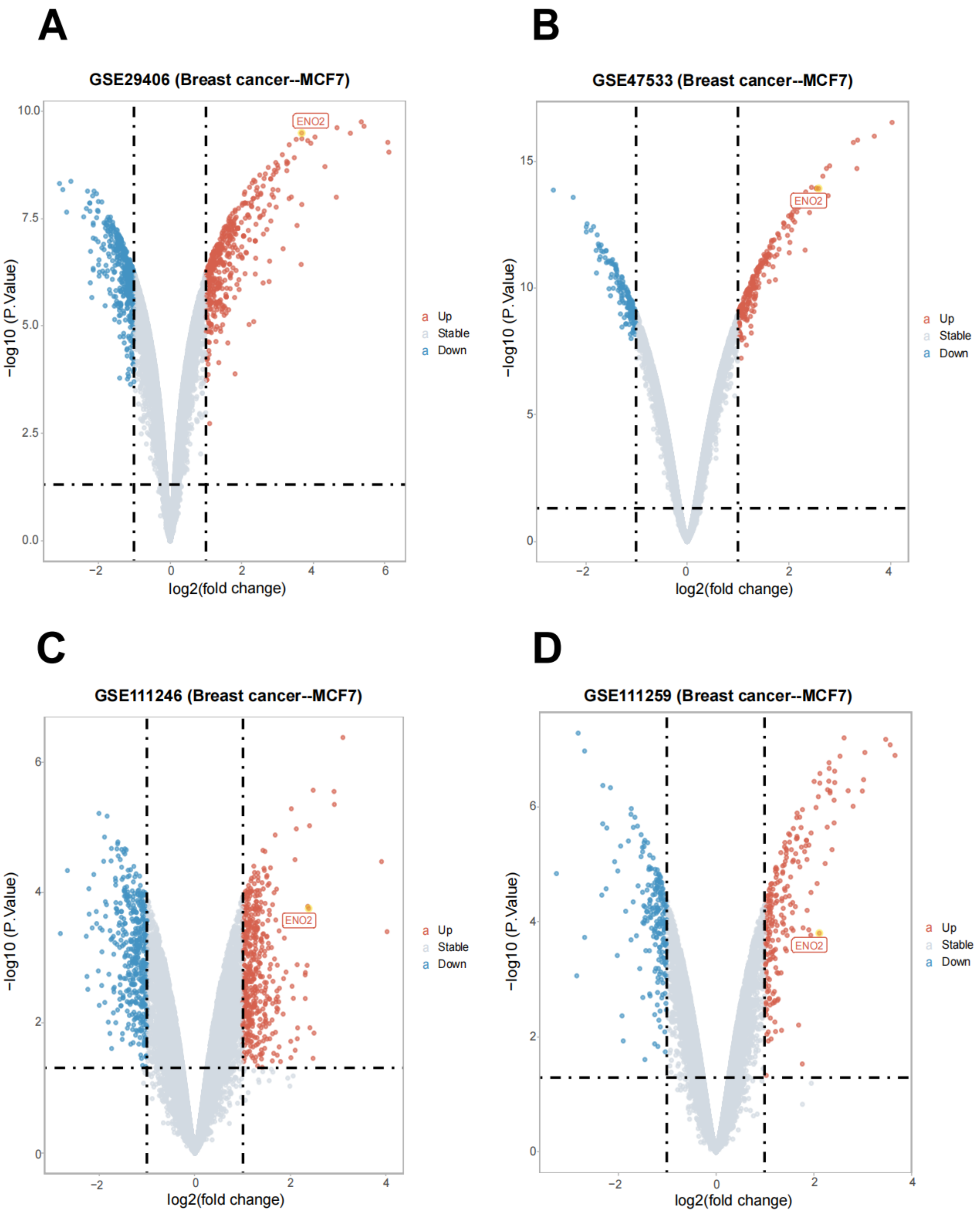

### Figure S2

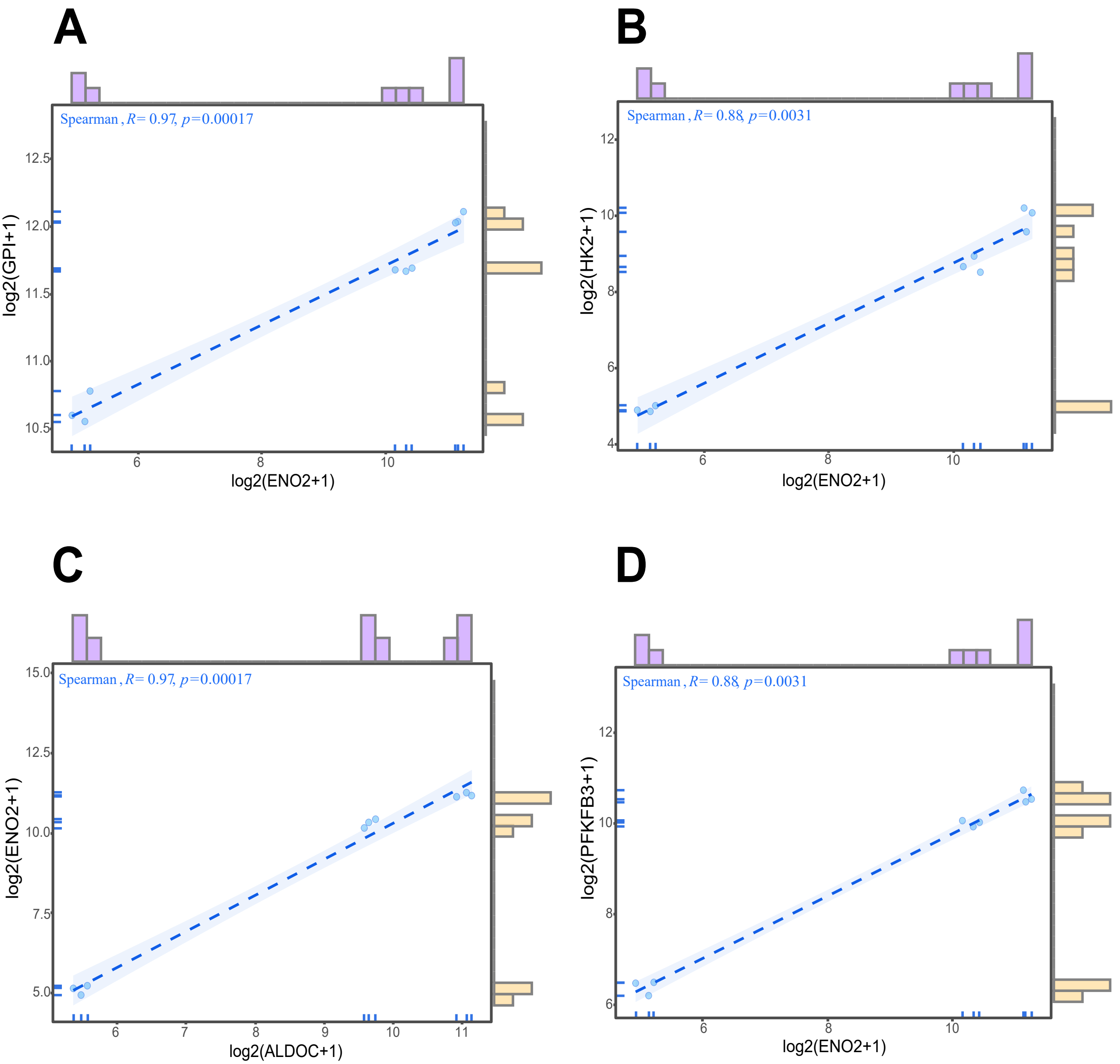
