## Supplementary material for "THER: Integrative Web Tool for Tumor Hypoxia Exploration and Research": Figure S3

A

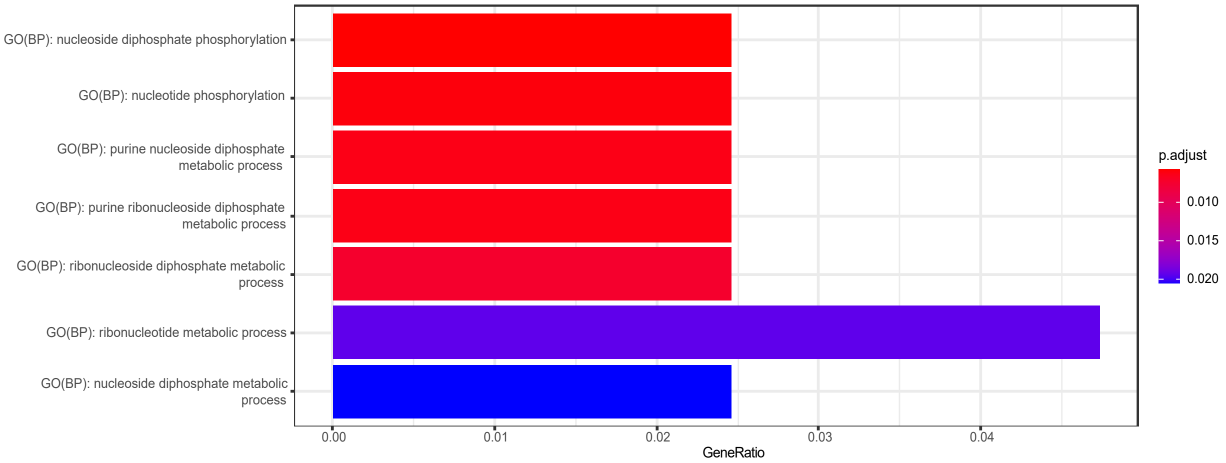


B

| | Description | GeneRatio | BgRatio | pvalue | p.adjust | qvalue | Count | geneID |
|---|---|---|---|---|---|---|---|---|
| GO:0006165 | GO(BP): nucleoside diphosphate phosphorylation | 13/528 | 132.00000 | 8.65e-05 | 5.77e-03 | 5.35e-03 | 13 | ENO2/DDIT4/ALDOC/HK2/PFKFB4/PFKFB3/PGM1/RANBP2/PFKP/TIGAR/HK1/GPI/NUP153 |
| GO:0046939 | GO(BP): nucleotide phosphorylation | 13/528 | 133.00000 | 9.34e-05 | 6.04e-03 | 5.60e-03 | 13 | ENO2/DDIT4/ALDOC/HK2/PFKFB4/PFKFB3/PGM1/RANBP2/PFKP/TIGAR/HK1/GPI/NUP153 |
| GO:0009135 | GO(BP): purine nucleoside diphosphate metabolic process | 13/528 | 135.00000 | 1.09e-04 | 6.35e-03 | 5.89e-03 | 13 | ENO2/DDIT4/ALDOC/HK2/PFKFB4/PFKFB3/PGM1/RANBP2/PFKP/TIGAR/HK1/GPI/NUP153 |
| GO:0009179 | GO(BP): purine ribonucleoside diphosphate metabolic process | 13/528 | 135.00000 | 1.09e-04 | 6.35e-03 | 5.89e-03 | 13 | ENO2/DDIT4/ALDOC/HK2/PFKFB4/PFKFB3/PGM1/RANBP2/PFKP/TIGAR/HK1/GPI/NUP153 |
| GO:0009185 | GO(BP): ribonucleoside diphosphate metabolic process | 13/528 | 138.00000 | 1.36e-04 | 7.42e-03 | 6.88e-03 | 13 | ENO2/DDIT4/ALDOC/HK2/PFKFB4/PFKFB3/PGM1/RANBP2/PFKP/TIGAR/HK1/GPI/NUP153 |
| GO:0009259 | GO(BP): ribonucleotide metabolic process | 25/528 | 425.00000 | 4.19e-04 | 1.93e-02 | 1.79e-02 | 25 | ENO2/DDIT4/ALDOC/HK2/PFKFB4/PFKFB3/PDK1/PGM1/BPNT1/UMPS/H19/CTPS1/BPNT2/SCD/ELOVL2/CROT/NT5C2/ATPSCKMT/RANBP2/PFKP/PDK3/TIGAR/HK1/GPI/NUP153 |
| GO:0009132 | GO(BP): nucleoside diphosphate metabolic process | 13/528 | 156.00000 | 4.55e-04 | 2.06e-02 | 1.91e-02 | 13 | ENO2/DDIT4/ALDOC/HK2/PFKFB4/PFKFB3/PGM1/RANBP2/PFKP/TIGAR/HK1/GPI/NUP153 |
